# Visuospatial working memory is cortically enabled through veridical, categorical and semantic representations

**DOI:** 10.1101/2025.08.08.669067

**Authors:** Joana Pereira Seabra, Andreea-Maria Gui, Vivien Chopurian, Alessandra S Souza, Carsten Allefeld, Thomas B Christophel

## Abstract

A single visual stimulus can elicit multiple concurrent representations throughout the cortex. We show that during a visual working memory task several cortical regions utilize categorical, semantic, and spatial representational formats to maintain visual stimuli in a robust fashion. We assessed the nature of orientation representations in an fMRI dataset of 40 participants performing an orientation memory task using multivariate encoding modelling. Our results show that orientation representations across the cortex form a gradient of abstraction, with more veridical representations in sensory areas and more abstract, categorical codes in anterior areas. We use cross-condition modelling to demonstrate shared neural codes between orientation and congruent verbal and location stimuli, used at different points during the trial. These findings provide evidence for a distributed account of working memory storage, where memory representations, distinct in nature and content, are present concurrently throughout the cortex.

## Introduction

The ability to accurately perceive and hold a stimulus in our mind for short periods of time is a key component of human cognition, enabling goal-directed behavior. These active, *working* memories are believed to rely on multiple cortical representations, diverse in nature and content (Christophel et al., 2017; Fuster, 1997). How do the contents of these representations differ from one another throughout the cortex, and why do we keep multiple simultaneous representations of single stimulus?

Visual working memory has been found to rely on numerous cortical areas, including sections of visual, parietal and prefrontal cortices (Christophel et al., 2017; Ester et al., 2015; Harrison & Tong, 2009). In nonhuman primates, both striate and extrastriate visual areas carry information about remembered stimuli (Chelazzi et al., 2001; Van Kerkoerle et al., 2017). Spatial information held in working memory has been found to be represented in parietal (Constantinidis & Steinmetz, 1996) and prefrontal areas (Funahashi et al., 1993), including the frontal eye fields (Armstrong et al., 2009). Prefrontal neurons exhibit stimulus-selective activity during memory in single-cell recordings for features like the frequency of a stimulus (Romo et al., 1999), its numerosity (Nieder et al., 2002), color (Buschman et al., 2011), and motion (Zaksas & Pasternak, 2006). In humans, multivariate pattern analysis (MVPA) has provided a more comprehensive understanding of the contents of cortical representations by making use of the patterned responses of cortical regions (Cox & Savoy, 2003; Haxby et al., 2013; Haynes & Rees, 2005; Kamitani & Tong, 2005). Using MVPA and fMRI, activity patterns representing remembered orientations and locations could be identified and decoded in early visual areas, IPS and sPCS (Christophel et al., 2018; Ester et al., 2015; Harrison & Tong, 2009; Sprague et al., 2016), regions similar to those evidenced by primate literature to hold spatial representations (Armstrong et al., 2009; Constantinidis & Steinmetz, 1996). These findings point to the existence of multiple concurrent working memory representations.

Holding multiple identical representations of the same stimulus would be costly and redundant. Alternatively, different cortical representations could hold the same stimulus in different yet complementary formats ranging from veridical to abstract, or from sensory to goal-dependent (Christophel et al., 2017; Fuster, 1997). Specifically, primary visual cortex would hold more veridical, fine grained memory traces, while representations from extrastriate visual cortex, to parietal cortex and frontal cortex, would increasingly be more categorical or abstract (Chunharas et al., 2024; Yan et al., 2023). These categorical representations are a likely cause of categorical biases in recall behavior which are more pronounced after extended delays (Bae, 2021; Bae et al., 2015; Balikou et al., 2015; Pereira Seabra et al., 2025; Romo et al., 1997). Verbalization of visual stimuli has been shown to increase the accuracy of recall (Overkott & Souza, 2022, 2023; Souza et al., 2021). Even when no instruction to verbalize is given, subjects frequently report verbalization throughout the delay (Chopurian et al., 2024; Pereira Seabra et al., 2025). Furthermore, patterns of labeling behavior are predictive of categorical biases (Bae et al., 2015; Pereira Seabra et al., 2025) as well as cortical categorical representations (Yan et al., 2023). Thus, a subset of cortical representations is likely to reflect this verbalization behavior. But not all abstractions of a stimulus need to be verbal or categorical. Cortical representations of remembered orientation and location stimuli largely overlap across the dorsal visual stream (Ester et al., 2015; Kwak & Curtis, 2022; Rademaker et al., 2019; Sprague et al., 2016), and orientation and location memory share behavioral biases and labeling patterns (Bae, 2021; Pereira Seabra et al., 2025). This suggests that these two forms of visual working memory rely on shared cortical resources.

The same stimulus can be represented by multiple regions across the cortical sheet and prior evidence suggests that these different regions correspond to different features, aspects or representations of the stimulus. What is detected of these representations is never fully static, they appear to wax and wane, and sometimes seemingly disappear (Christophel et al., 2018; Fuster, 1973; Iamshchinina et al., 2021; Lewis-Peacock et al., 2012; Murray et al., 2017). Sometimes, even the neuronal code underlying a memory representation - the pattern of selective neuronal responses representing a stimulus - appears to change over time such that generalization between time points is reduced or absent (Emrich et al., 2013; Li & Curtis, 2023; Stokes et al., 2013). This dynamic coding is often described as a fundamental computational feature (Stroud et al., 2024), a rapid series of states (Stokes et al., 2013), or a trajectory in feature space (Meyers, 2018). Working memory, however, is not a passive store but an active workspace where items can change in meaningful ways (Baddeley, 1992). For example, representations of a stimulus before and after a mental rotation in the same cortical regions use shared cortical codes to represent changing content (Albers et al., 2013; Christophel et al., 2015).

Here, we systematically assess the representational content of cortical working memory stores to provide a more comprehensive delineation of the formats of storage underlying visual working memory. We first ask whether multiple representations of a single visuo-spatial item held in memory represent these same contents differently. If so, which ones reflect more abstract, categorical codes, and which rely on more veridical, continuous codes? Secondly, we explore the relation between working memory representations of orientation stimuli and congruent verbal stimuli by assessing whether their representations exhibit a common semantic code. Lastly, we inquire whether the memory of a simpler visuo-spatial stimulus (locations) could elicit representations akin to those of an orientation to understand if orientation working memory relies, at least partially, in a simpler visual format. We find a set of cortical representations using categorical, semantic, and spatial codes stably shared across tasks.

## Results

### Visual and anterior regions form a gradient of abstraction

We investigated the representational nature of visual working memory representations focusing on data recorded while subjects (N = 40) performed a single-item orientation recall task. Subjects memorized a centrally presented orientation stimulus (Fig. 1A) and recalled this orientation after an extended delay (15.2 s) by rotating a probe until it matched the remembered content. Critically, we sampled the orientation space in a detailed fashion using 24 orientation stimuli that were evenly distributed and presented with equal frequency. This allowed us to map fine-grained representational changes between different areas and across the unfolding of the extended memory delay. Subjects were able to recall the remembered orientations reliably (absolute error: *M* = 7.74°, *SEM* ± 0.13). Consistently with previous findings (Balikou et al., 2015; Essock, 1980; Huttenlocher et al., 1991; Pereira Seabra et al., 2025), we found that absolute error for cardinal orientations was significantly smaller than that of non-cardinal orientations (t_39_ = −7.98, p < .001), and that near-cardinal stimuli were recalled more diagonally (i.e., as shifted away from the cardinals) than other non-cardinal orientations (t_39_ = 6.34, p < .001). Both results are indicative of categorization processes during visual working memory.

**Figure 1:**
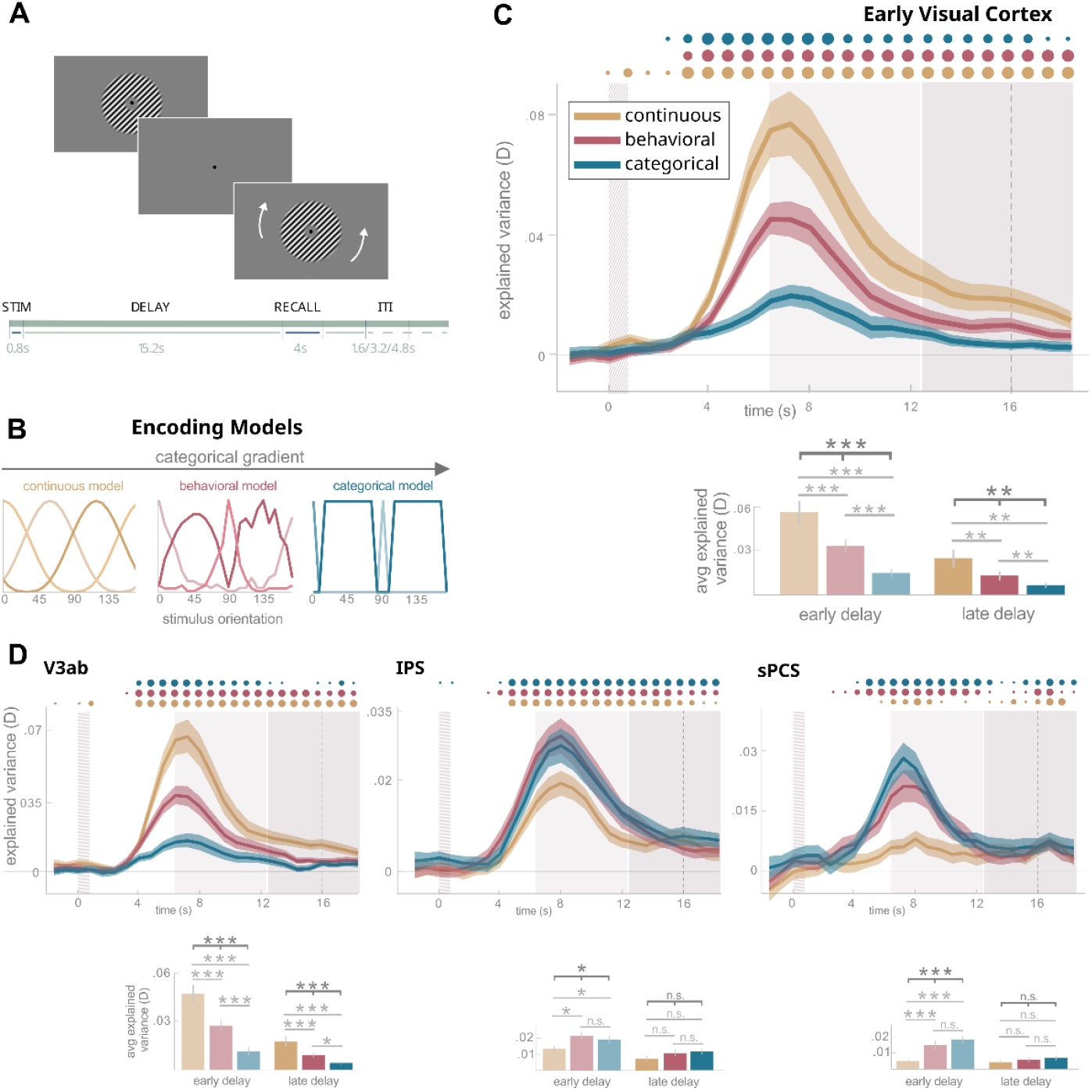
Encoding modelling shows preference for continuous model in posterior regions, and for categorical and behavioral-based models in anterior regions. **A**. Orientation memory task. Participants were instructed to memorize the stimulus and, after a delay, rotate a probe to match its initial orientation. The stimulus was presented for 0.8s, followed by a 15.2s delay and a 4s recall period. Inter-trial interval was randomly set to 1.6, 3.2 or 4.8s. **B**. We used three encoding models to model representations of memorized content in fMRI data, organized along a categorical gradient from more continuous to more categorical. **C-D**. Timeseries plots of each model’s performance explaining stimulus-specific variance in fMRI data from the orientation task, averaged across all participants and trials for each ROI. Ribbons indicate SEM. The first gray area corresponds to the stimulus presentation period, the second light-gray area represents the early delay period, and the third darked-gray one represents the late delay. The barplots also show each model’s performance in two partitions of the delay period, now averaged across all timepoints included in each partition (i.e. early or late, 6.4s to 12.8s and 12.8s to 18.4s, respectively). Above the plot, dots varying in size indicate the significance of one-sided paired t-tests evaluating the difference between explained variance values obtained by each model and baseline (0), in each TR (every 0.8s).One-way ANOVAs were used to assess interaction effects between the three models during early (all *p* < .05) and late delays (for both EVC and V3b *p* < .01, for IPS and sPCS *p* > .05) and paired t-tests were used to compare between pairs of models (for EVC and V3b all *p* < .05, see figure for IPS and sPCS).

To determine the representational content of cortical regions carrying information about working memory contents, we first asked whether the information is stored in a more continuous or categorical format. For this, we compared whether variance in stimulus-specific activity representing memorized contents in visual, parietal and frontal regions is best explained by a continuous, an intermediary (behavior-based), or a strictly categorical encoding model (see Figure 1A). Using three-regressor models (see Methods for details), in early visual cortex and V3ab, during the early delay, the continuous model outperformed both the behavioral (EVC: *t*_39_ = 3.95, *p* < .001; V3ab: *t*_39_ = 4.32, *p* < .001) and the categorical models (EVC: *t*_39_ = 4.66, *p* < .001; V3ab: *t*_39_ = 5.67, *p* < .001). The same pattern of results was found during the late delay (all *p* < .01). In both regions and time-periods, the intermediary, behavioral model also outperforms the categorical model (all *p* < .05). In more anterior regions, in contrast, the behavioral and categorical models explain the underlying data better during the early and late delay, compared to posterior areas. In the early delay, both the behavioral and categorical models outperform the continuous model in IPS (behavioral: *t*_39_ = −3.11, *p* = .01; categorical: *t*_40_ = −2.62, *p* = .025) and sPCS (behavioral: *t*_39_ = −3.84, *p* < .001; categorical: *t*_39_ = −5.17, *p* < .001). While the intermediary model was preferred in IPS, and the categorical model explained most variance in sPCS, no significant differences were found when comparing these models directly (IPS: *t*_39_ = 0.64, *p* = .52 ; sPCS: *t*_39_ = −1.47, *p* = .15). In the late delay, no significant differences between the three models were found in either anterior region (all *p* > .58). Anterior and posterior regions appear to form a gradient of abstraction where stimulus-specific activity in early visual areas is best explained by continuous encoding models, and anterior working memory stores are best explained by intermediary or fully abstracted categorical models.

### Semantic, verbal representations partially explain stimulus-selectivity in visual cortex

Given the evidence for the presence of multiple representational formats across the cortex, we then asked if some of these could take a verbal, semantic form. To investigate this, we asked subjects to perform a separate auditory and visual word 1-back task using spatial language (see Figure 3A). We trained a multivariate encoding model to distinguish activity recorded during the visual and auditory processing of spatial words (‘horizontal’, ‘left’, ‘vertical’, and ‘right’; see Methods section for a more detailed explanation) used during orientation rehearsal in prior work (Pereira Seabra et al., 2025). We then asked whether the same model could be used to distinguish orientation corresponding to these words (modeled using a 4-regressor continuous model, see Fig. 2B). Using this cross-decoding approach, we found that orientation-selective responses could be predicted using this four-word semantic model in early visual cortex (*t*_39_ = 3.67, *p* < .001) and in V3ab (*t*_39_ = 2.89, *p* = .007) during the early delay, but not during the late delay (both *p* > .1). In anterior areas, explained variance resulting from the cross-decoding analysis did not significantly differ from baseline in neither early nor late delays (all *p* > .1). Using the behavior or the strictly categorical models for similar analysis resulted in a qualitatively similar pattern of results. This directly shows that during encoding and the early stages of the delay, fMRI activity patterns in posterior areas resemble activity during the processing of spatial language.

**Figure 2:**
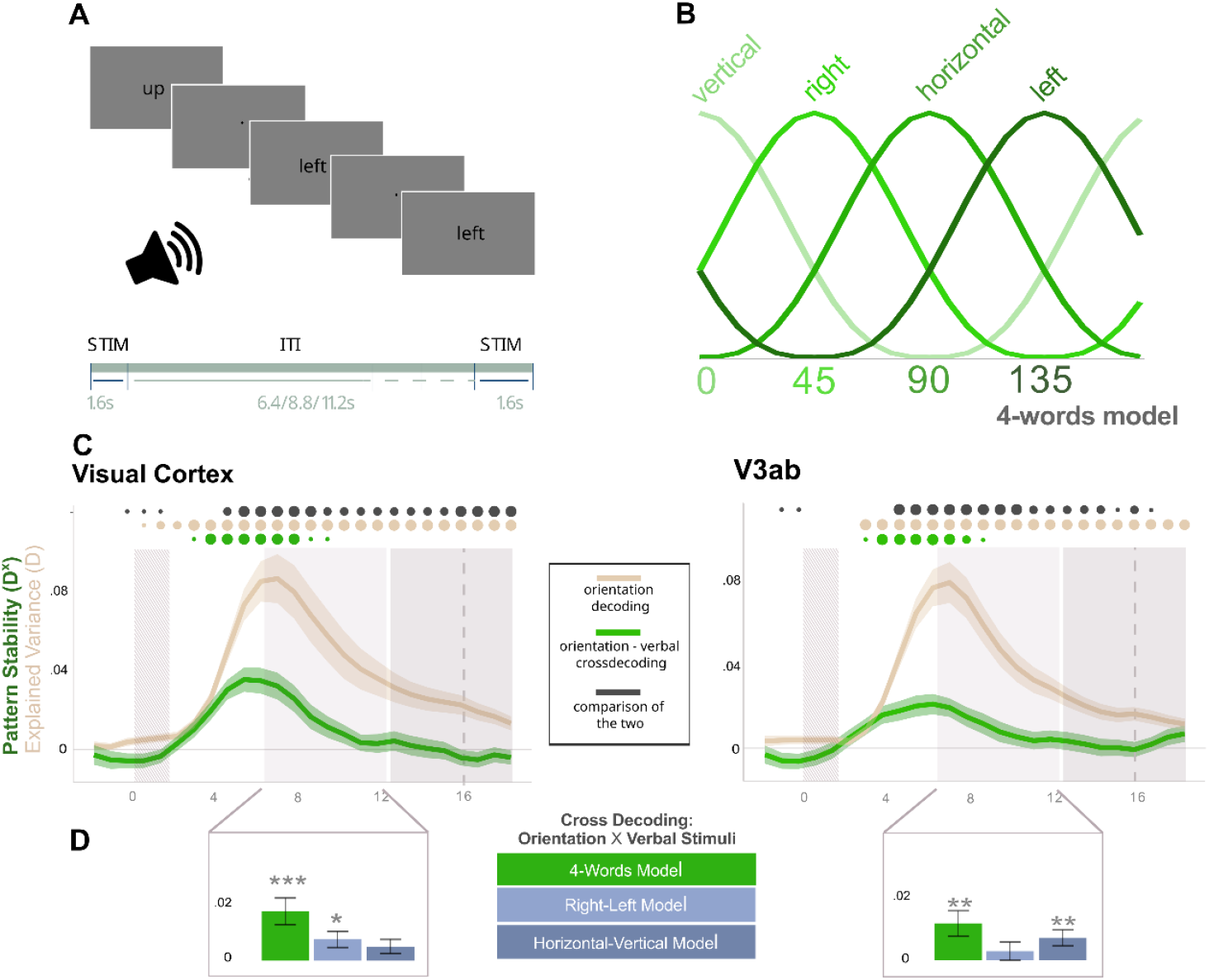
Orientation representations during working memory share activity patterns with data from a verbal task. **A**. Verbal 1-back task example trial. Participants were instructed to memorize the word and, after a delay, indicate whether the next word was the same by pressing a button. Words were presented both visually and auditorily in each trial. The stimulus was presented for 2s, followed by a 6.4, 8.8 or 11.2s delay. **B**. The 4-words continuous model used for the training data. Each regressor was based in a word for which its usage significantly predicted recall error for orientations in a previous behavioral study (Pereira Seabra et al., 2025). The regressor’s are equidistant, and each center was defined based on the most common stimulus rotation positions associated with that word in the behavioral study. **C**. Time series plots of cross decoding results, in green, with a model trained on neural data recorded during word presentation in the verbal task and tested on orientation task data. For reference, in beige, results from the continuous model trained and tested on orientation data from Fig 1. Above the plot, dots varying in size indicate the significance of one-sided paired t-tests evaluating the difference between explained variance values obtained by each model and baseline (0), in each TR (every 0.8s). The dark grey dots represent results from a comparison between the two models (orientation-orientation, and verbal-orientation) also with one-sided paired t-tests. **D**. Explained variance averaged across all time points in the early delay in each ROI for each verbal model, and comparison against baseline with a one-sided paired t-tests (in visual cortex, for the 4-words model *p* <. 001 and for the right-left model *p* = .013; in V3ab, for the 4-words model and horizontal-vertical model, both *p* <.01).

**Figure 3:**
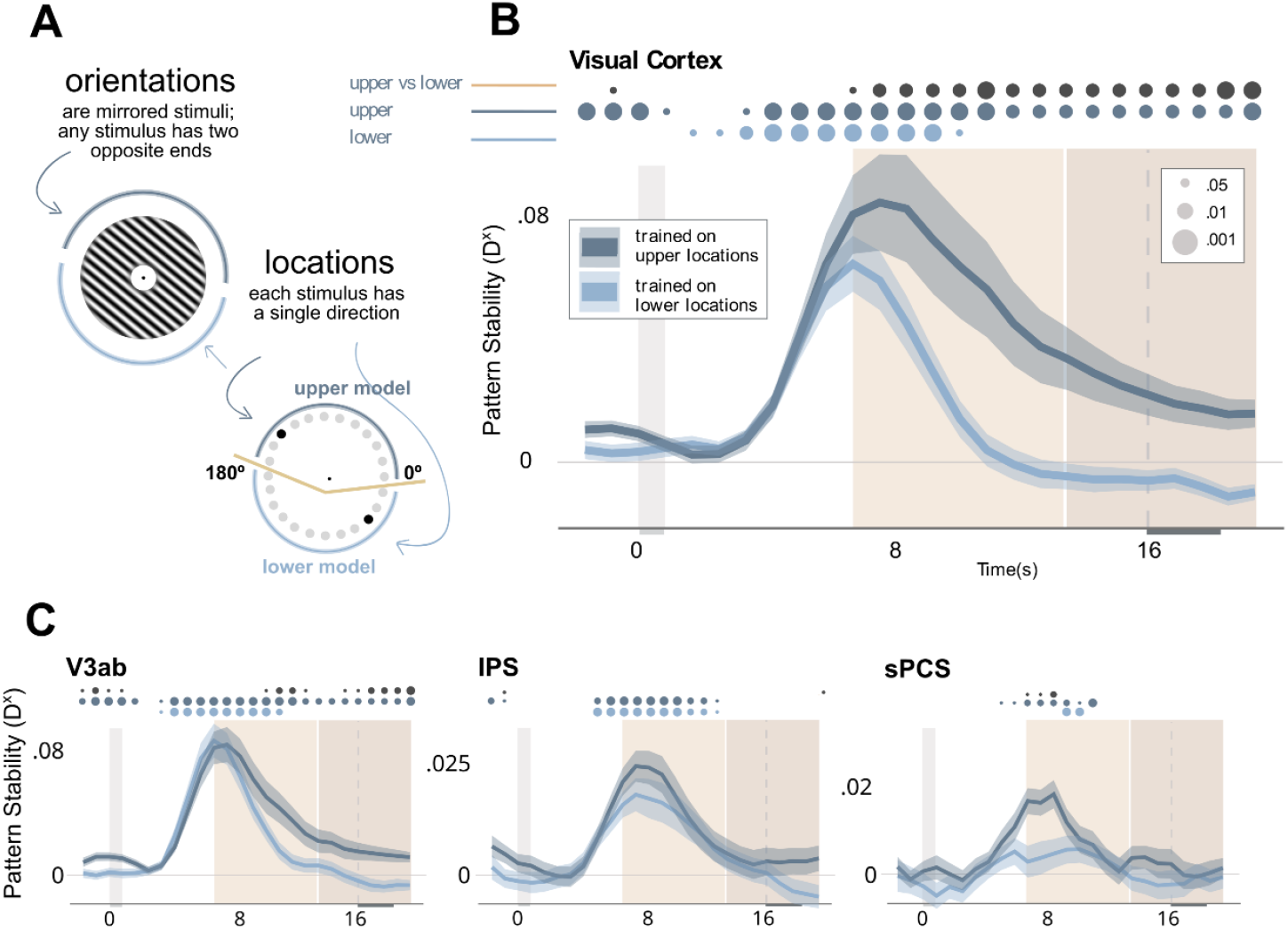
Orientation and location representations during working memory share an abstracted representational space. **A**. Orientation and location stimuli used in the task, and the division of location stimuli used to create the upper and lower training models. **B - C**. Time series plot of cross-decoding results in early visual cortex, V3ab, IPS and sPCs. The dark blue line represents the mean explained variance and respective standard error of the mean between orientations and upper locations, while the light blue one represents the lower locations model. The early partition of the delay is represented in the plot by the sand-colored box, while the darker sand color represents the late delay. Above the plot, in the corresponding shades of blue, dots indicate the significance of one-sided paired t-tests evaluating the difference between explained variance values obtained by each model and baseline (0), in each TR (every 0.8s). The dark grey dots represent results from a comparison between the two models, also with one-sided paired t-tests.

To understand whether this semantic cross-decoding relied exclusively on cardinal or diagonal orientations, we divided the initial continuous 4-regressor model for the verbal data into 2 models. The first model featured basis functions centered on cardinal stimuli (0° and 90°), corresponding to the words ‘horizontal’ and ‘vertical’, and the second model centered on diagonal stimuli (45° and 135°), corresponding to ‘right’ and ‘left’. In the early delay, the ‘right-left’ model performs above chance in early visual cortex (*t*_39_ = 2.62, *p* = .013), and the ‘horizontal-vertical’ model in V3ab (*t*_39_ = 2.82, *p* = .008), suggesting that while explained variance appears to be reduced for these less-informed models both model variants can result in robust cross-decoding.

### Orientation memory relies on simplified spatial representations

Lastly, we inquired whether orientation representations relied on a simplified spatial code. To investigate this, subjects performed a location recall task (identical to the orientation task, but with location stimuli). We cross-decoded between neural responses to location and orientation stimuli during working memory by training models on the former and testing them on the latter. This analysis assumes that, during orientation working memory tasks, subjects simplify the Gabor stimulus and only retain the endpoint(s) of a central line of the Gabor patch. As every orientation stimulus has two endpoints, we performed the analysis twice, once with a model including only the upper locations, and the other with only the lower locations, while all orientations remained the same in both comparisons (see Methods for details).

In posterior areas, both the upper and lower location models were able to explain orientation-selective activity during the early delay (all *p* < .001), but only the upper model was able to do it during the late delay (EVC: *t*_39_ = 3.4, *p* = .001; V3ab: *t*_39_ = 3.5, *p* = .001). In both posterior areas the upper model had a significant advantage over the lower model during the late delay (EVC : *t*_39_ = 3.3, *p* = .002, V3ab: *t*_39_ = 2.1, *p* = .043), but only during the early delay in primary visual cortex (*t*_39_ = 2.7, *p* = .01, V3ab: *t*_39_ = .5, *p* = .63). In anterior areas, the upper model was able to cross-decode orientations during the early delay (IPS: *t*_39_ = 6.4, *p* < .001; sPCS: *t*_39_ = 5.25, *p* < .001), but the lower model could only be confirmed to do so in IPS (*t*_39_ = 4.7, *p* < .001, sPCS: (*t*_39_ = 1.6, *p* = .13). During the late delay in anterior areas, we found no cross-decoding results that significantly differed from baseline (all *p* > .05). We found an upper model advantage in anterior regions only in sPCS during the early delay (*t*_39_ =2.7,*p* = .009, all other *p* > .1), where the lower model did not significantly differ from baseline. Notably, sPCS is where this advantage emerges first in the timeline (see Fig. 4C) and it is unclear whether this preference becomes more difficult to detect in the late delay as decodable information drops, or whether sPCS represents both spatial locations for redundancy during these later stages of the trial. Thus, all regions appear to employ a shared neural code to retain orientation information, simplifying angle information to one or two spatial locations.

## Discussion

In this study, we set out to investigate the nature of distinct neural representations elicited by a single oriented grating held in working memory. We find that regions holding stimulus-specific information about remembered orientations form a posterior to anterior gradient from veridical, continuous representations to exceedingly more abstracted and categorical representations. A model trained on activity during language processing of spatial words shows that a subset of stimulus-specific information in visual cortices can be explained by semantic representations. Across all regions, orientation working memory is, at least partially, carried by simplified spatial codes, with either one or both endpoints of a given orientation being represented at different time points. We observe dynamic changes in the relative contribution of most of these memory representations over time.

This representational gradient of abstraction has long been postulated (Christophel et al., 2017; Fuster, 1997) and is generally consistent with the gradual processing of perceptual inputs (Dumoulin & Wandell, 2007; Tootell et al., 1998; Wager & Smith, 2003). In prior work, evidence for this gradient during working memory but not during perceptual tasks has been found for color (Yan et al., 2023) and orientations (Chunharas et al., 2024). Here, we provide direct evidence that representations ranging from veridical to categorical, including verbal, are present throughout the cortex during working memory. Biases increase when comparing perceptual tasks to memory ones (Bae et al., 2015), but also tend to increase with time within memory at intermediate and high memory load (Zhou et al., 2022).These behavioral changes are directly predictable based on our neural data. This seems to be further consistent with subjective experience, as participants report they categorize and label visual stimuli, and their free labeling behavior predicts categorical biases (Bae et al., 2015; Pereira Seabra et al., 2025). Thus, this work suggests that behavioral strategies and biases can be informative of the underlying neural processes. But not all biases or internalized simplifications and abstractions are known. A focus on behavior and subjective reports is thus critical to further our understanding of neural representations.

We used labeling data from a previous study (Pereira Seabra et al., 2025) to define categorization, but also investigated semantic processing directly by training encoding models on data obtained during spatial language processing and testing these models on orientation working memory data. We find this kind of generalization between codes from drastically different tasks in the early visual cortex and V3ab, early in the delay. This suggests visual working memory relies partly on verbal cortical representations particularly during encoding. So while categorization has been found to relate to the verbalization of visual stimuli (Grill-Spector & Kanwisher, 2005; Zettersten & Lupyan, 2020), so much so that several categorization models account for a verbal component (Minda et al., 2024), our semantic cross-decoding results identify semantic representations during working memory that are regionally separate from categorical working memory codes. While these results seems counterintuitive, behavioral evidence suggests that asking subjects to verbally represent visual stimuli increases the retention of continuous representations (Souza et al., 2021; Souza & Skóra, 2017). Notably, categorization does not require verbalization (Minda et al., 2024) and categorization effects have been found previously in visual areas (Ester et al., 2020; Henderson et al., 2025). More generally, word representations have been found to extend into visual and motor cortices (Huth et al., 2016; Mitchell et al., 2008) and visual processing has been shown to be a necessary element of spatial language processing (Coventry et al., 2013). Hence, semantic cortical processing of visual working memory contents might be limited or at least focused on the encoding phase and not overlap with categorization effects. Whether other verbal-like representations in more anterior regions exist during visual working memory in forms that are inaccessible to our cross-decoding methods is a question for future research.

A third form of abstraction is spatial simplification. When training models on neural responses to location stimuli and testing them on responses to orientations, our cross-decoding results are consistent with evidence for line-like spatial abstractions of orientations and moving dot stimuli (Kwak & Curtis, 2022). Data presented here suggests that subjects might be capable of simplifying an orientation even further, representing it by retaining just one location on its edge. This abstraction strategy likely reduces effort and is driven by attentional biases towards the upper visual hemifield (Langley & McBeath, 2023; Zito et al., 2016) as evidenced by the model trained on upper locations outperforming the lower locations one. Upper attentional biases also affect labeling behavior (Pereira Seabra et al., 2025). These biases may originate in anterior regions (Leavitt et al., 2018) and propagate to posterior areas as the trial unfolds.

Working memory is inherently understood not as a passive store but as an adaptive, changing workspace (Baddeley, 1992; Christophel et al., 2017). This capability to adaptively change what is being represented is demonstrated by task related changes in the localization of storage (Lee & Baker, 2016), and in changes in cortical represented contents due to mental rotation (Albers et al., 2013; Christophel et al., 2015). Thus, malleability might be a central feature of cortical working memory representations (Kiyonaga & Serences, forthcoming). Here, we provide evidence for the presence of multiple, separable neural representations elicited by low-level visuospatial stimuli, and the nature of these representations, ranging from veridical to abstract. Representations leaning on less veridical codes can hold different types of abstract information, such as visual simplifications or verbal labels.

## Methods

### Participants

Forty-five participants (ages 19–36 years; mean age: 26, SEM ± 0.75) gave informed consent and took part in the study. All were fluent in English, right-handed, with normal or corrected-to-normal vision. Each participant completed two 90-minute sessions of MRI, at least one day apart. Ethical approval was granted by the Ethics Committee of the Institute of Psychology of Humboldt-Universität zu Berlin. Out of the initial 45 participants, 40 were included in the final sample. Three subjects did not finish the full experiment and data for two subjects were missing because of data corruption or loss.

### Experimental Design

#### Orientation and Location Memory Task

Across the first and second session, participants completed a total of 16 runs of a delayed estimation task: 8 of these runs featured orientation stimuli, and another 8 runs featured location stimuli. Both tasks are identical. Participants were shown a single stimulus for 0.8s followed by a 15.2s delay after which a probe appeared for participants to match its rotation to the original stimulus (see Fig. 1A). The task was performed by rotating the probe with buttons the subjects had in each hand, and once they were confident that the rotation of the probe was as similar as possible to the initial sample, pressing another button to log their answer within 2.4s. Before the next trial, there was an interval of either 1.6s, 3.2s or 4.8s. In each run, participants performed a total of 24 trials, each with a unique rotation of the stimulus.

#### Verbal Task

In the second session, after subjects had finished the orientation and location memory task, participants completed a word 1-back task using spatial language. In each trial, a spatial word stimulus (e.g., ‘left’) was presented twice (sequentially) as a typed word in the center of the screen for 0.6s each time, simultaneously with an audio recording of the corresponding word (interstimulus interval = 0.8s). This was followed by 6.4s, 8.8s, or 11.2s of intertrial interval. Participants were instructed to press a button every time the words repeated in two sequential trials. There were a total of 4 runs of 42 trials each (6 per word). Until they were instructed to do it inside the scanner, subjects were unaware of the nature of this task or the stimuli used.

#### Stimuli

The experiment included three types of stimuli: orientations, locations and words. The orientation stimuli were high-contrast grayscale circular Gabor patches (422 pixel radius) with varying phase, constant spatial frequency, and edge smoothing of 60 pixels. A fixation (10 pixels) dot was presented in a central annulus (60 pixel radius). We used 24 orientations as sample stimuli (from 0° to 172.5°; 7.5° apart). The location stimulus was a single dot (20 pixels) presented on an imaginary circle at a distance of 424 pixels from fixation. The location of the dot on the imaginary circle was defined using polar coordinates. We used 24 samples (from 0° to 345°, 15° apart). Please note that due to the point symmetry inherent in grating stimuli, the stimulus spaces are either limited to 180° (orientations) or 360° (locations), resulting in different stimulus spacing in the two tasks. For the verbal task, stimuli were selected among the most common spatial language terms used to describe orientation and location stimuli in a free naming task (Pereira Seabra et al., 2025). The selected terms used as stimuli were: ‘horizontal’, ‘vertical’, ‘diagonal’, ‘left’, ‘right’, ‘up’ and ‘down’. Audio stimuli were generated using an online text-to-speech generator (https://speechgen.io) with a neutral-toned female-sounding voice with a generic US-English accent (option ‘Joanna’ on the website).

### Data Acquisition

All the fMRI recordings were obtained from the same MRI scanner (3T Siemens Tim Trio) with the Syngo MR XA30 operating system, located in Freie Universität Berlin. VisuaStim projection equipment was used for stimulation and response collection was performed using MR-compatible button boxes. The experimental display projection was 48cm wide and 27.2cm tall, and viewing distance was 100 to 110cm. 711 functional images were recorded in each of the 16 runs with an orientation and location memory task, and 561 functional images for each of the 4 verbal task runs (72contiguous slices, TR = 800ms, TE = 37ms, voxel size = 2mm^3^, flip angle = 52°, slice gap = 0mm, FOV = 208mm). After the 16 memory task runs, before the verbal task runs, a T1-weighted MPRAGE structural image was recorded (176 contiguous slices, TR = 1900ms, TE = 2.52ms, Voxel size = 1mm^3^, flip angle = 9°, slice gap = 1mm, FOV = 256mm).

### Imaging Data

#### Preprocessing

The imaging data were preprocessed with Matlab R2022a (The MathWorks Inc., 2022) using SPM12 (Friston et al., 1994) and CvCrossManova toolboox v0.0.0 (released November 2023; Allefeld & Haynes, 2014). Functional and anatomical images were converted from the originally acquired format (DICOM) into NIFTI for compatibility with the software. All functional images were realigned to match the positioning of the first one in order to correct for head movements. The anatomical image was coregistered to this first functional image and segmented to estimate normalization parameters. The anatomical data were also normalized (voxel resolution = 2mm^3^) and spatially smoothed (smoothing kernel = 5mm^3^). Behavioral data were analyzed using R version 4.1.1 (R Core Team, 2021) and R Studio (RStudio Team, 2020).

#### Regions of Interest

We investigated four regions of interest known to represent orientations and locations during working memory (Rademaker et al., 2019; Sprague et al., 2016): early visual cortex (V1-V3), V3ab, IPS and sPCS. For this, we used probabilistic maps of human visual topography available online (Wang et al., 2015). All maps included both left and right hemisphere locations, and the early visual cortex included dorsal and ventral regions. The early visual cortex map was created by combining regions V1 to V3, V3ab was comprised of V3a and V3b, IPS included IPS1 to IPS6. These combined ROIs were warped into single subject space by applying inverse normalization parameters obtained during preprocessing. We narrowed down the regions of these maps using activation-based masks generated using stimulation models with standard HRF-regressors for the presentation of each kind of stimulus (i.e. orientation, location and verbal). We selected the 1000 most active voxels based on t-maps contrasting all stimuli for each modality in each subject, over a threshold of 10% probability of belonging to the ROI (Chopurian et al., 2024; Yan et al., 2023). For cross-modality encoding models we selected voxels in similar fashion, but selected the 1000 most active voxels of both modalities combined.

### Analyses

#### Encoding Modelling for Orientation Data

We tested the representational content of these regions of interest using encoding modelling (Allefeld & Haynes, 2014; Brouwer & Heeger, 2009; Yan et al., 2023). An encoding model describes the selective neuronal response of a set of voxels to a feature (here, orientation) using a set of regressors called basis functions. These basis functions can be seen quantifying the tuning of a voxel analogous to tuning curves of single cells (Roth et al., 2012; Wang et al., 2003). The shape of these basis functions thus carries information about how narrow or wide, and how constant or variable the selectivity of a voxel is.

In a first step, three encoding models with three basis functions each were used to assess the nature of the neural data in the orientation memory tasks in early visual cortex, V3ab, IPS and sPCS - a continuous, a behavior-based and a categorical model (Fig. 1B). The first was a bell-shaped model, where the shape of each basis function is continuous (similar to a normal distribution) and the width remains equal across basis functions. This type of continuous model has been used throughout prior work as an approximator for a sensory-like encoding model (Brouwer & Heeger, 2009; Chunharas et al., 2024; Ester et al., 2015; Yan et al., 2023). In the current study, parameter *x* represents the stimulus space and *μ* the center of each basis function (60, 120 or 180 degrees):

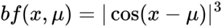

The behavioral model was based on orientation naming data from a previous study from our group (Pereira Seabra et al., 2025, data available at https://osf.io/n27jd/). We generated 3 basis functions, each based on how frequently each of the most recurringly used words (‘horizontal’, ‘vertical’ or ‘diagonal’) was used to describe each stimulus. These basis functions were normalized. The categorical model is a boxcar-shaped model with regressors of 2 different widths, specifically 1 or 18 stimuli. The two basis functions with a single stimulus correspond to stimuli rotated at 0 or 90 degrees, while the third basis function includes the remaining range of stimuli from 15 to 75 and 105 to 165 degrees that are frequently referred to by participants as ‘diagonal’. For each basis function, the average response across all stimuli (e.g., all orientations) was subtracted from every response, turning the regressor into a de facto weighted contrast between one set of stimuli versus another set.

We tested how much each model explained the neural response during the orientation task using CvMANOVA (Yan et al., 2023; for a predecessor, see Allefeld & Haynes, 2014). CvMANOVA is a type of multivariate pattern analysis that estimates how well an encoding model explains variance between neural activity patterns for different stimuli. This measure of explained variance, pattern distinctness (D), is cross-validated across separate sets of training and test data. If the data contains no stimulus-specific variance, D varies around 0. Otherwise, D expresses a ratio of explained variance divided by unexplained variance. The analysis was performed on the selected voxels of each ROI (see section ‘Regions of Interest’) from 1.6 seconds before stimulus presentation, until 18.4 seconds after stimulus presentation. This time period is equivalent to 26 TRs of 0.8 seconds, for which we used 26 finite impulse response (FIR) regressors. For each three-basis-function model, the design matrix modeled 703 scans per run with 78 regressors (3 basis functions x 26 FIR bins). We then estimated the variance jointly explained by each set of three basis functions by estimating model parameters for these basis functions using seven out of eight runs and testing on the remaining run. This procedure was repeated until every run was used as test data, once (8-fold cross-validation). We used a regularization parameter of Lambda = 0.9 for estimation. This approach to encoding modelling was extensively validated using simulated data (Christophel et al., 2024).

#### Cross-decoding Orientation and Verbal Data

We then assessed whether it was possible to cross-decode orientation stimuli from neural data attained during the verbal task from EVC, V3ab, IPS and sPCS. Neural responses to the presentation of four words (i.e. ‘horizontal’, ‘vertical’, ‘left’ and ‘right’) were modelled using four conventional HRF-regressors. The words were selected based on orientation naming data from a previous study from our group (Pereira Seabra et al., 2025). We here selected four words instead of three since ‘diagonal’ seems to correspond to a much wider area than ‘horizontal’ or ‘vertical’. In order to use equidistant regressors with the same width, the four words above seemed like the most sensible choice. The resulting parameter estimates were tested for generalization with a continuous model comprised of 4 basis functions for the orientation data (see above). For this cross-decoding analysis, CvCrossManova was used (method not yet fully published, but see Allefeld, 2024a, 2024b; Iamshchinina et al., 2021) to estimate pattern stability (D^x^), i.e. the shared explained variance between the two data sets. Each word regressor was paired with a continuous regressor centered around the orientation that the word usually refers to (specifically, 0° = vertical, 45° = right, 90° = horizontal, 135° = right). Explained variance for this cross-condition model was estimated using procedures described above, but here for each validation fold, we selected three out of the four verbal task runs and tested on one of the eight orientation runs. Given the experiment was only comprised of four verbal runs, each of the four different validation sets was repeated once. Orientations scans are modelled like in the previous analysis (703 scans, 104 regressors) and the 553 scans in each verbal run are modeled by 4 regressors.

To test whether this verbal cross-decoding relied exclusively on the distinction between cardinal and non-cardinal, left and right, or horizontal and vertical orientations (and their corresponding word representations), we also tested two alternative models including only the words ‘left’ and ‘right’, or only ‘horizontal’ and ‘vertical’ and the corresponding continuous regressors.

#### Cross-decoding Orientations from Location Data

Finally, we assessed whether it was possible to cross-decode orientations and locations in EVC, V3ab, IPS and sPCS with CvCrossManova as well. To easily map orientations and locations onto each other, a model comprising 6 boxcar regressors –binning 4 orientations togethers, at a time - was used to model the data. While locations can cover the entire 360 degrees of a circular space, a mirrored stimulus like orientations can only encompass 180 degrees. To address this difference, we built two training models for the location data, one with upper locations - 0 to 165 degrees - and another with lower locations −180 to 345 degrees (Fig. 4A) and used both individually to cross-decode the corresponding orientation stimuli. The analyses were validated, with each training set consisting of 7 location runs, and the testing set of 1 orientation run mimicking the cross-validation structure from the initial orientation-only analyses. Unlike in the other analyses, we contrasted the basis functions in pairs (e.g. bf1 against bf2, bf2 against bf3, and so on), as the boxcar regressors did not inherently contrast different stimuli against each other. The analyses followed the same procedure to the one described above, except there were 156 regressors (6 x 26 FIR bins) modelling the 703 scans.

#### Statistical testing

In all analyses, explained variance for each model was estimated for each of the 26 timepoints individually, allowing us to investigate the changes in explained variance over time. We used one-sided one-sample t-test to test explained variance (D) against chance (D = 0) and pairwise paired t-tests were used to compare pairs of models in both the early and late delays.

#### Behavioral Data Analysis

We assessed behavioral accuracy by computing the trial wise absolute circular error (remembered – recalled stimulus). Cardinal biases were analyzed by comparing absolute recall error differences between cardinal (0°, 90°, 180°, 270°) and non-cardinal stimuli (all other stimuli) with one-sided paired t-tests. Directional biases were assessed by testing near-cardinal stimuli (up to 15° away in either direction) against stimuli far from the cardinals (22.5° to 67.5°). Relative recall error was used in these analyses, i.e. averaged recall error after correcting for most frequent error direction in each stimulus. The far and near-cardinal stimuli groups were compared using one-sided paired t-tests.

## Supplements

## Author Contributions

JPS, TC and ASS conceptualized and designed research. JPS, VC and TC programmed the experiment. JPS acquired and analyzed data, with the support of TC, CA and AMG. CA provided valuable assistance in the implementation of CvCrossManova. AMG provided valuable insights from prior analyses related to the project. JPS wrote the original draft. TC, ASS, VC, CA, and AMG reviewed and edited the manuscript.

## Ethics and Consent

This project was performed in accordance with the Declaration of Helsinki and received ethical approval by the internal review board of the Institute for Psychology of the Humboldt-Universität zu Berlin (IRB number: 2022–04). Participants signed an informed consent form and their identity was anonymized.

## Acknowledgements

The authors would like to thank Damla Cifci, Rita Bertani and Zhiqi Kang for their support during fMRI data collection.

## Funding Information

TBC received funding by a DFG Emmy Noether Research Group grant CH 1674/2-1.

## Competing Interests

The authors have no competing interests to declare.

## Notes

### Competing Interest Statement

The authors have declared no competing interest.

### Summary of Updates

Figures were reuploaded with increased resolution

